# Determination of host cell proteins constituting the molecular microenvironment of coronavirus replicase complexes by proximity-labeling

**DOI:** 10.1101/417907

**Authors:** V’kovski Philip, Gerber Markus, Kelly Jenna, Pfaender Stephanie, Ebert Nadine, Braga Lagache Sophie, Simillion Cedric, Portmann Jasmine, Stalder Hanspeter, Gaschen Véronique, Bruggmann Remy, Stoffel Michael, Heller Manfred, Dijkman Ronald, Thiel Volker

**Affiliations:** Institute of Virology and Immunology IVI, Bern, Switzerland; Department of Infectious Diseases and Pathobiology, Vetsuisse Faculty, University of Bern, Bern, Switzerland; Graduate School for Biomedical Science, University of Bern, Bern, Switzerland; Interfaculty Bioinformatics Unit and SIB Swiss Institute of Bioinformatics, University of Bern, Switzerland; Mass Spectrometry and Proteomics Core Facility, Department for BioMedical Research (DBMR), University of Bern, Switzerland; Department of Clinical Research, University of Bern, Switzerland; Division Veterinary Anatomy, Vetsuisse Faculty, University of Bern, Bern, Switzerland

**Keywords:** RNA virus, Coronavirus, replication complex, virus-host interaction, replication compartments, proximity-labeling, viral replicase molecular microenvironment.

## Abstract

Positive-sense RNA viruses hijack intracellular membranes that provide niches for viral RNA synthesis and a platform for interactions with host proteins. However, little is known about host factors at the interface between replicase complexes and the host cytoplasm. We engineered a biotin ligase into a coronaviral replication/transcription complex (RTC) and identified >500 host proteins constituting the RTC microenvironment. siRNA-silencing of each RTC-proximal host factor demonstrated importance of vesicular trafficking pathways, ubiquitin-dependent and autophagy-related processes, and translation initiation factors. Notably, detection of translation initiation factors at the RTC was instrumental to visualize and demonstrate active translation proximal to replication complexes of several coronaviruses.

Collectively, we establish a spatial link between viral RNA synthesis and diverse host factors of unprecedented breadth. Our data may serve as a paradigm for other positive-strand RNA viruses and provide a starting point for a comprehensive analysis of critical virus-host interactions that represent targets for therapeutic intervention.

## Introduction

Positive-strand RNA viruses replicate at membranous structures that accommodate the viral replication complex and facilitate RNA synthesis in the cytosol of infected host cells (1-5). Rewiring host endomembranes is hypothesized to provide a privileged microenvironment physically separated from the cytosol, thereby ensuring adequate concentrations of macromolecules for viral RNA synthesis, preventing recognition of replication intermediates such as double-stranded RNA (dsRNA) by cytosolic innate immune receptors (6, 7), and providing a platform that facilitates molecular interactions with host cell proteins.

Ultrastructural studies have reported the origin, nature, and extent of membrane modifications induced by coronaviruses (order *Nidovirales*, family *Coronaviridae*), which materialize as an ER-derived network of interconnected double-membrane vesicles (DMVs) and convoluted membranes (CM) in perinuclear regions of infected cells to which the viral replication/transcription complex (RTC) is anchored (4, 8, 9). The RTC is generated by translation of the genomic RNA into two large polyproteins that are extensively auto-proteolytically processed by viral proteases to give rise to 16 processing end-products, termed non-structural proteins (nsps) 1-16. Nsp1 is rapidly cleaved from the polyproteins and not considered an integral component of the coronaviral RTC, but interferes with host cell translation by inducing degradation of cellular mRNAs (10-12). Although it has not yet been formally demonstrated, the remaining nsps (2-16) are thought to comprise the RTC and harbor multiple enzymes and functions, such as de-ubiquitination, proteases, helicase, polymerase, exo-and endonuclease, and N7-and 2’O-methyltransferases (13-18). Many of these functions have been studied using reverse genetic approaches, which revealed their importance in virus-host interactions (19-23). In most cases phenotypes were described via loss-of-function mutagenesis, however, in the context of virus infection, the specific interactions of RTC components with host cell factors remain largely unknown.

A number of individual host cell proteins have been shown to impact coronavirus replication by using various screening methods, such as genome-wide siRNA, kinome, and yeast-two-hybrid screens (24-28). Likewise, genome-wide CRISPR-based screens have been applied to other positive-stranded RNA viruses, such as flaviviruses, and identified critical host proteins required for replication (29, 30). Some of these proteins were described in the context of distinct ER processes, such as N-linked glycosylation, ER-associated protein degradation (ERAD), and signal peptide insertion and processing. Although individual proteins identified by these screens may interact with viral replication complexes, they likely constitute only a small fraction of the global replicase microenvironment.

To capture the full breadth of host cell proteins and cellular pathways that are spatially associated with viral RTCs, we employed a proximity-based labeling approach involving a promiscuous *E. coli*-derived biotin ligase (BirA_R118G_). BirA_R118G_ biotinylates proximal (<10 nm) proteins in live cells without disrupting intracellular membranes or protein complexes, and hence, does not rely on high affinity protein-protein interactions, but is able to permanently tag transient interactions (31). Covalent protein biotinylation allows stringent lysis and washing conditions during affinity purification and subsequent mass spectrometric identification of captured factors. By engineering a recombinant MHV harboring BirA_R118G_ as an integral component of the RTC we identified >500 host proteins reflecting the molecular microenvironment of MHV replication structures. siRNA-mediated silencing of each of these factors highlighted, amongst others, the functional importance of vesicular ER-Golgi apparatus trafficking pathways, ubiquitin-dependent and autophagy-related catabolic processes, and translation initiation factors. Importantly, the detection of active translation in close proximity to the viral RTC highlighted the critical involvement of translation initiation factors during coronavirus replication. Collectively, the determination of the coronavirus RTC-associated microenvironment provides a functional and spatial link between conserved host cell processes and viral RNA synthesis, and highlights potential targets for the development of novel antiviral agents.

## Results

### Engineering the BirA_R118_-biotin ligase into the MHV replicase transcriptase complex

To insert the promiscuous biotin ligase BirA_R118G_ as an integral subunit of the MHV RTC, we used a vaccinia virus-based reverse genetic system (32, 33) to generate a recombinant MHV harboring a myc-tagged BirA_R118G_ fused to nsp2. This strategy was recently employed by Freeman *et al.* for a fusion of green fluorescent protein (GFP) with nsp2 (34). MHV-BirA_R118G_-nsp2 retained the cleavage site between nsp1 and BirA_R118G_, while a deleted cleavage site between BirA_R118G_ and nsp2 ensured the expression of a BirA_R118G_-nsp2 fusion protein (Fig. 1a).

**Figure 1.**
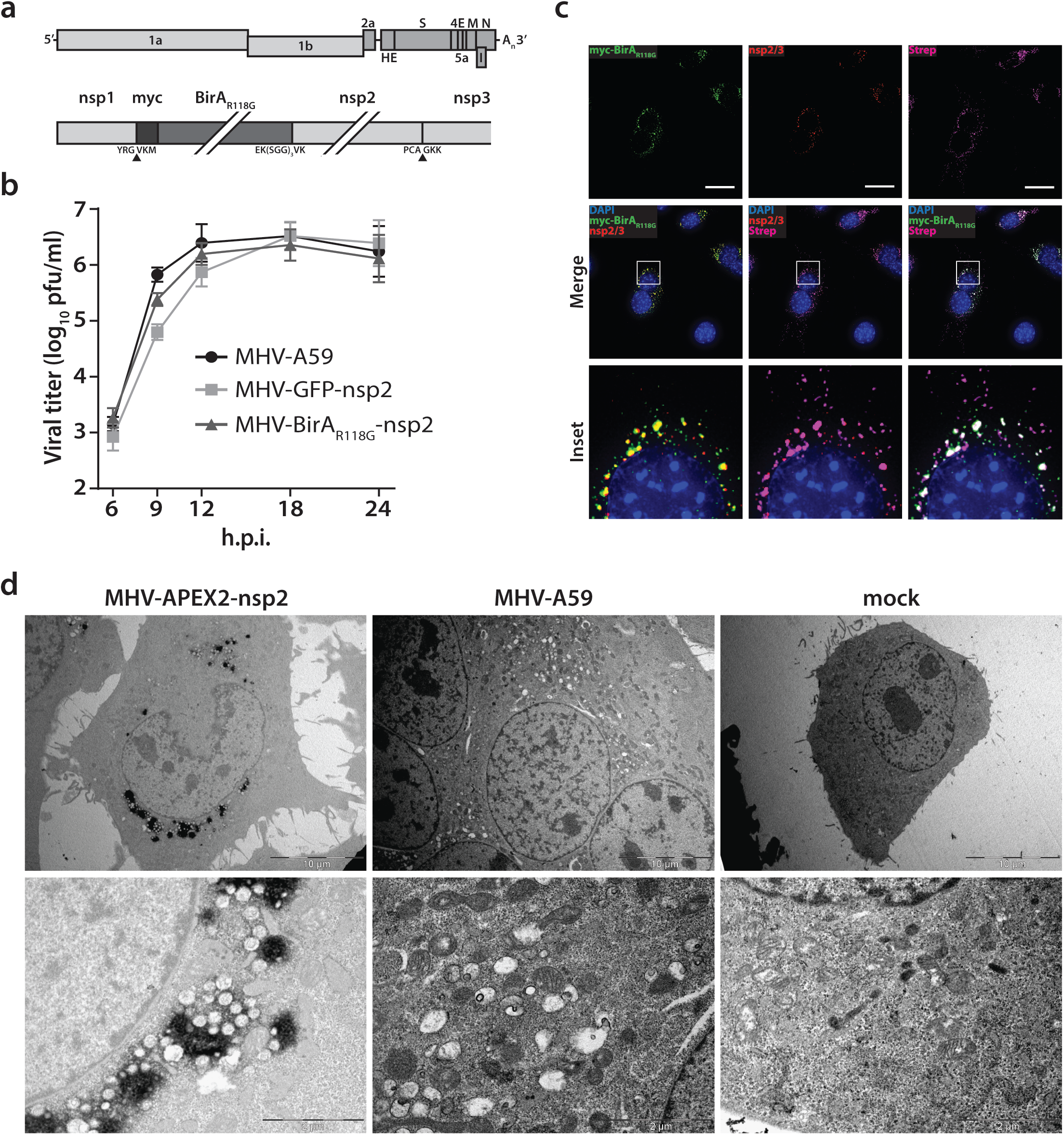
Characterization of the recombinant MHV-BirA_R118G_-nsp2. (a) Genome organization of recombinant MHV-BirA_R118G_-nsp2. The positive-sense RNA genome of MHV contains a 5’ cap and a 3’ poly(A) tail. ORF1a and ORF1b encode the viral replication and transcription complex (nsp1-16). myc-BirA_R118G_ was inserted as an N-terminal fusion with nsp2 within ORF1a. The cleavage site between nsp1 and myc-BirA_R118G_ was retained (black arrow) while a deleted cleavage site between BirA_R118G_ and nsp2 ensured the release of a BirA_R118G_-nsp2 fusion protein from the pp1a polyprotein. The cleavage site between nsp2 and nsp3 was also retained. (b) Viral replication kinetics of recombinant MHV-BirA_R118G_-nsp2 were compared to wild-type MHV-A59 and recombinant MHV-GFP-nsp2. Murine L929 fibroblasts were infected at a multiplicity of infection (MOI) of 1 plaque forming unit (pfu) per cell. Viral supernatants were collected at the indicated time points, titrated by plaque assay and expressed in pfu per ml. Data points represent the mean and SEM of three independent experiments, each performed in quadruplicate. (c) Immunofluorescence analysis of MHV-BirA_R118G_-nsp2-mediated biotinylation of RTC-proximal factors. L929 cells were infected with MHV-BirA_R118G_-nsp2 (MOI=1) in medium supplemented with 67µM biotin. Cells were fixed 15 hours post infection (h.p.i.) and processed for immunofluorescence analysis with antibodies directed against the BirA_R118G_ (anti-myc), the viral replicase (anti-nsp2/3) and biotinylated factors (streptavidin). Nuclei are counterstained with DAPI. Z-projection of deconvolved z-stacks acquired with a DeltaVision Elite High-Resolution imaging system are shown. Scale bars: 20 µm. (d) Ultrastructural analysis of MHV-APEX2-nsp2 infection. L929 cells were infected with MHV-APEX2-nsp2 and MHV-A59 (MOI=2), or mock infected. At 10 h.p.i., cells were fixed, stained with DAB and processed for electron microscopy investigations. Representative low and high magnifications are displayed.

MHV-BirA_R118G_-nsp2 replicated to comparable peak titers and replication kinetics as the parental wild-type MHV-A59 (Fig. 1b). MHV-EGFP-nsp2, which was constructed in parallel and contained the coding sequence of EGFP (34) instead of BirA_R118G_, was used as a control and also reached wild-type virus peak titers, with slightly reduced viral titers at 9 hours post-infection (h.p.i.) compared to MHV-A59 and MHV-BirA_R118G_-nsp2 (Fig. 1b).

To confirm the accommodation of BirA_R118G_ within the viral RTC, MHV-A59-, MHV-BirA_R118G_-nsp2-, and mock-infected L929 fibroblasts were visualized using indirect immunofluorescence microscopy. BirA_R118G_-nsp2 remained strongly associated with the MHV RTC, as indicated by the co-localization of BirA_R118G_-nsp2 with established markers of the MHV replicase, such as nsp2/3 and nsp8 (Fig. 1c; Supplemental Fig. S1). This observation corroborates previous studies demonstrating that nsp2, although not required for viral RNA synthesis, co-localizes with other nsps of the coronaviral RTC (35-37). Importantly, by supplementing the culture medium with biotin, we could readily detect biotinylated proteins with fluorophore-coupled streptavidin that appeared close to the MHV RTC in MHV-BirA_R118G_-nsp2-infected cells, demonstrating efficient proximity-dependent biotinylation of RTC-proximal host factors (Fig. 1c; Supplemental Fig. S1). Furthermore, to define the localization of the nsp2 fusion protein at the ultrastructural level, we replaced the BirA_R118G_ biotin ligase with the APEX2 ascorbate peroxidase to generate recombinant MHV-APEX2-nsp2. APEX2 mediates the catalysis of 3,3’-diaminobenzidine (DAB) into an insoluble polymer that can be readily observed by electron microscopy (38). As shown in figure 1d, APEX2-catalized DAB polymer deposition was readily detectable at characteristic coronavirus replication compartments, such as DMVs and CM, categorically demonstrating that the nsp2 fusion proteins localize to known sites of coronavirus replication (4, 8).

Importantly, our collective results establish that the recombinant MHV-BirA_R118G_-nsp2 replicates with comparable kinetics to wild-type MHV-A59, expresses a functional BirA_R118G_ biotin ligase that is tightly associated with the MHV RTC, and that biotinylated, RTC-proximal proteins can be readily detected in MHV-BirA_R118G_-nsp2 infected cells.

### Determination of the coronavirus RTC-proximal proteome

To further demonstrate the efficiency and specificity of BirA_R118G_-mediated biotinylation we assessed, by western blot analysis, fractions of biotinylated proteins derived from MHV-A59-, MHV-BirA_R118G_-nsp2-, or non-infected cells that were grown with or without the addition of biotin (Fig. 2a,b). A characteristic pattern of endogenously biotinylated proteins was observed under all conditions where no exogenous biotin was added to the culture medium (Fig. 2b). The same pattern was detectable in non-infected and wild-type MHV-A59-infected cells when the culture medium was supplemented with biotin, suggesting that the addition of biotin in the absence of the BirA_R118G_ biotin ligase does not recognizably change the fraction of endogenously biotinylated proteins. In contrast, we observed a greatly increased fraction of biotinylated proteins in lysates derived from MHV-BirA_R118G_-nsp2-infected cells treated with biotin. This result demonstrates that virus-mediated expression of the BirA_R118G_ biotin ligase results in efficient biotinylation when biotin is added to the culture medium. Moreover, we could readily affinity purify, enrich, and recover the fraction of biotinylated proteins under stringent denaturing lysis and washing conditions by using streptavidin-coupled magnetic beads (Fig. 2b).

**Figure 2.**
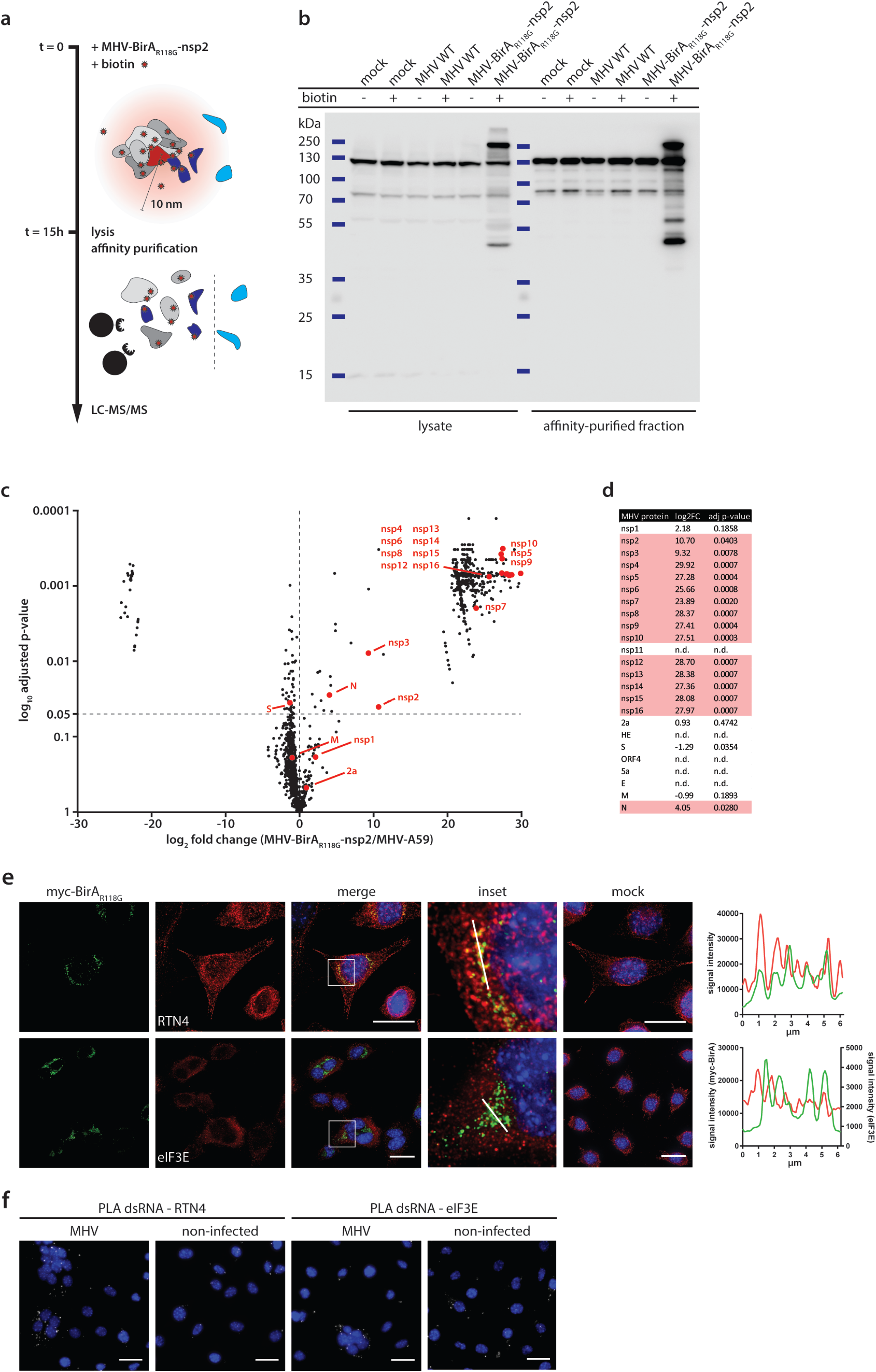
Determination of the coronavirus RTC-proximal proteome. (a) Schematic overview of the BirA_R118G_-mediated proximity biotinylation assay using MHV-BirA_R118G_-nsp2. (b) Western blot analysis of MHV-BirA_R118G_-nsp2-infected L929 cells. L929 cells were infected with MHV-BirA_R118G_-nsp2, MHV-A59 (parental wild-type strain) or non-infected in medium with and without supplementation of 67 µM biotin. Cells were lysed 15 h.p.i. and biotinylated factors were subjected to affinity purification using streptavidin-coupled magnetic beads. Total cell lysates and affinity-purified fractions were separated by SDS-PAGE and analysed by western blot probed with horse radish peroxidase (HRP)-coupled Streptavidin. (c) Host and viral factors identified by LC-MS/MS. 4*107 L929 cells were infected with MHV-BirA_R118G_-nsp2 or MHV-A59 in medium supplemented with 67µM biotin. 15 h.p.i., lysates were affinity purified and LC-MS/MS was performed from in-gel digested samples. MS identification of biotinylated proteins was performed in three independent biological replicates. Spectral interpretation was performed against a Mus musculus and MHV database and log_2_-transformed LFQ levels (x-axis) were used to determine significant differences in protein enrichment between sample groups (Student’s T-test, y-axis). Identified cellular proteins are displayed as black dots, MHV proteins are highlighted in red (nsp: non-structural protein, N: nucleocapsid, S: spike, M: membrane, 2a: accessory protein 2a). (d) Summary of viral proteins identified by LC-MS/MS. nsp2-10, nsp12-16, and nucleocapsid were significantly enriched in fractions derived from MHV-BirAR118G-nsp2-infected cells whereas nsp1, nsp11, structural proteins spike (S), envelope (E) and membrane proteins (M) as well as all accessory proteins (NS2a, HE, ORF4, ORF5a) were either not significantly enriched or not detected. (e,f) Immunofluorescence analysis of RTC-proximal cellular factors. L929 cells were seeded on coverslips, infected with MHV-BirA_R118G_-nsp2 (e) or MHV-A59 (f), fixed at 9 h.p.i. and processed for immunofluorescence using anti-myc, anti-RTN4 and anti-eIF3E antibodies (e) or anti-dsRNA, anti-RTN4 and anti-eIF3E antibodies (f). Secondary fluorophore-coupled antibodies were used to detect the viral replicase and endogenous levels of RTN4 and eIF3E (e). Proximity ligations were performed using Duolink In Situ detection reagents (f). Nuclei are counterstained with DAPI. Z-projection of deconvolved z-stacks acquired with a DeltaVision Elite High-Resolution imaging system are shown. Intensity profiles highlighted in the magnified regions are shown. Scale bars: 20 µm.

Affinity purified proteins derived from biotin-treated MHV-A59-and MHV-BirA_R118G_-nsp2-infected cells were subjected to mass spectrometric analysis (n=3). Liquid chromatography tandem-mass spectrometry (LC-MS/MS) was performed from in-gel digested samples and log-transformed label free quantification (LFQ) levels were used to compare protein enrichment between samples (Fig. 2c). Overall, 1381 host proteins were identified, of which 513 were statistically significantly enriched in MHV-BirA_R118G_-nsp2-infected samples over MHV-A59-infected samples. These host proteins represent a comprehensive repertoire of RTC-proximal factors throughout MHV infection (Fig. 2c, table S1). Importantly, viral replicase gene products nsp2-10 and nsp12-16, and the nucleocapsid protein were significantly enriched in fractions derived from MHV-BirA_R118G_-nsp2-infected cells (Fig. 2c, d). This is in agreement with studies demonstrating co-localization and interactions amongst individual nsps, and with studies showing association of the nucleocapsid protein with the coronavirus RTC (8, 39-41). It also highlights the specificity and effectiveness of the labeling approach in live cells and is the first experimental evidence showing that collectively these viral nsps and the nucleocapsid (N) protein are subunits of the coronavirus RTC. Furthermore, these results corroborate previous reports that nsp1 is likely not an integral component of the coronavirus RTC (10-12, 42). Amongst the “not detected” or “not enriched” viral proteins are (i) nsp11, which is a short peptide of only 14 amino acids at the carboxyterminus of polyprotein 1a with a yet unassigned role or function in coronavirus replication, (ii) the structural proteins spike (S) protein, envelope (E) protein, and membrane (M) protein, which mainly localize to sites of viral assembly before being incorporated into newly-formed viral particles, and (iii) all accessory proteins (NS2a, HE, ORF4, ORF5a). Altogether, these results validate the proximity-dependent biotinylation approach and demonstrate the specific and exclusive labeling of MHV-RTC-associated proteins (Fig. 2d).

The BirA_R118G_ biotin ligase biotinylates proteins in its close proximity that must not necessarily have tight, prolonged, or direct interaction (31). Therefore, the identified RTC-proximal host proteins, recorded over the entire duration of the MHV replication cycle, likely include proteins that display a prolonged co-localization with the MHV RTC, proteins that may locate only transiently in close proximity to the RTC, and proteins of which only a minor fraction of the cellular pool may associate with the RTC. To this end, we assessed the localization of a limited number of host proteins from our candidate list in MHV-infected cells. Accordingly, we identified RTC-proximal host proteins displaying a pronounced co-localization with the MHV RTC, such as the ER protein reticulon 4 (rtn4; Fig. 2e), and host proteins where co-localization by indirect immunofluorescence microscopy was not readily detectable, such as the eukaryotic translation initiation factor 3E (eIF3E; Fig. 2e). However, in the latter case, a more sensitive detection technique, such as a proximity ligation assay that relies on proximity-dependent antibody-coupled DNA probe amplification (43), demonstrated proximity of eIF3E and dsRNA in MHV-infected cells (Fig. 2f).

Collectively, our results show that the approach of integrating a promiscuous biotin ligase as an integral subunit into a coronavirus RTC revealed a comprehensive list of host cell proteins that comprises the RTC microenvironment. The efficacy and specificity of our approach is best illustrated by the fact that we were able to identify all expected viral components of the MHV RTC, while other viral proteins, such as nsp1, structural proteins S, E, and M, and accessory proteins, were not amongst the significantly enriched proteins. Our data further suggest that the RTC microenvironment may be highly dynamic and likely also contains proteins that are only transiently present in the microenvironment or only comprise a sub-fraction of the cellular pool in close proximity to the MHV RTC.

### Functional classification of RTC-proximal host factors

To categorize functionally-related proteins from the list of RTC-proximal host proteins and identify enriched biological themes in the dataset, we performed a functional classification of RTC-proximal factors using Gene Ontology (GO) enrichment analysis. 86 GO biological process (BP) terms were significantly enriched in the dataset (p-value <0.05), of which 32 terms were highly significant (p-value <0.005) (Fig. 3a, Table S2). Additional analysis using AmiGO revealed that 25 of these 32 highly significant GO BP terms fell into 5 broad functional categories, namely cell adhesion, transport, cell organization, translation, and catabolic processes. To examine these categories further, identify important cellular pathways within them, and extract known functional associations among RTC-proximal host proteins, we performed STRING network analysis on the RTC-proximal proteins in each category (Fig. 3b, c, Fig S2).

**Figure 3.**
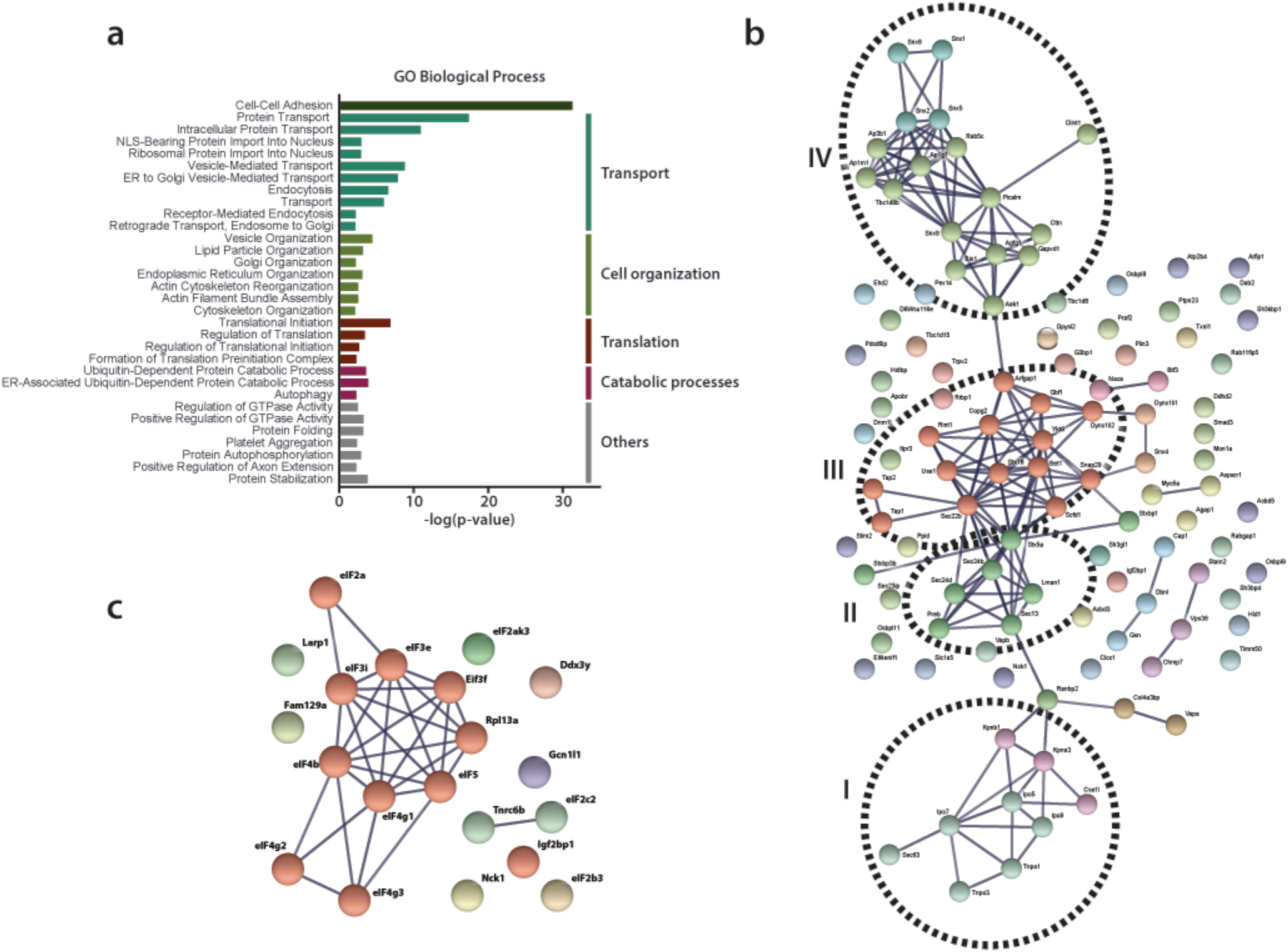
Functional classification of RTC-proximal host factors. (a) Gene Ontology enrichment analysis of RTC-proximal cellular factors. 32 terms were highly significant (p-value <0.005) and were assigned to 5 broad functional categories: cell-cell adhesion, transport, cell organization, translation, catabolic processes. (b-c) STRING protein interaction network analysis of the categories “transport” (b) and “translation” (c). The nodes represent RTC-proximal host proteins and the edges represent the interactions, either direct (physical) or indirect (functional), between two proteins in the network. Cellular proteins assigned to the “transport” category separated into 4 distinct interaction clusters. I: protein transport, II: COPII anterograde transport, III: COPI retrograde transport, IV: clathrin-mediated transport.

Despite “cell-cell adhesion” scoring high, it likely represents a typical limitation of gene annotation databases, where many genes play multiple roles in numerous pathways and processes. Accordingly, most genes assigned to the GO BP term cell-cell adhesion are also found in the other categories described below.

The category “transport” included protein trafficking and vesicular-mediated transport pathways and comprised the majority of RTC-proximal factors (Fig. 3a, b). Protein interaction network analysis, using STRING, revealed at least 4 distinct clusters of interacting factors within this category (Fig. 3b). Cluster I, protein transport, comprised nuclear transport receptors at nuclear pore complexes, such as importins and transportins. Interestingly, this cluster also contained Sec63, which is part of the Sec61 translocon (44) and has been implicated in protein translocation across ER membranes. The list of RTC-proximal factors also included signal recognition particles SRP54a and SRP68 (Table S2) proteins that promote the transfer of newly synthetized integral membrane proteins or secreted proteins across translocon complexes. Furthermore, the list contained Naca and BTF3, which prevent the translocation of non-secretory proteins towards the ER lumen (45, 46). Interestingly, genome-wide CRISPR screens have identified proteins involved in biosynthesis of membrane and secretory proteins as required for flavivirus replication (29, 30), suggesting similarities between flaviruses and coronaviruses concerning the requirement of ER-associated protein sorting complexes for viral replication.

Cluster II included vesicle components, tethers and SNARE (Soluble N-ethylmaleimide-sensitive-factor Attachment protein Receptor) proteins characteristic of the COPII-mediated ER-to-Golgi apparatus anterograde vesicular transport pathway whereas, cluster III contained components of the COPI-related retrograde Golgi-to-ER transport machinery. Moreover, Cluster IV was comprised of proteins that mediate clathrin-coated vesicle (endosomal) transport between the plasma membrane and the trans-Golgi network (TGN), which is also closely associated with the actin cytoskeleton. Together with sorting nexins, cluster IV components can be regarded as regulating late-Golgi trafficking events and interacting with the endosomal system.

Many of the cellular processes and host proteins assigned to “transport” (specifically in clusters II-IV) are also listed in the category “cell organization” (Fig. 3a, S2a). However, this category actually extends the importance of vesicular transport as it also contains factors involved in the architecture, organization, and homeostasis of the ER and Golgi apparatus, and the cytoskeleton-supporting these organelles. The prominent appearance of biological processes linked to protein and vesicular transport between the ER and both the cis-and trans-Golgi network, is in agreement with previous findings that have reported the relevance of the early secretory pathway for a number of RNA viruses, including coronaviruses (24, 26, 47-50).

Notably, a number of MHV RTC-proximal factors were part of the host translation machinery and assigned to category “translation” (Fig. 3a, c). We found enrichment of factors involved in the initiation of translation, particularly multiple subunits of eIF3 and eIF4 complexes, as well as eIF2, eIF5, the Ddx3y helicase, and the Elongation factor-like GTPase 1, which are required for the formation of 43S pre-initiation complexes, 48S initiation complexes, and the assembly of elongation-competent 80S ribosomes (51). The high degree of interaction between these subunits is suggestive of the presence of the entire translation initiation apparatus in close proximity to the viral RTC. The 60S ribosomal protein L13a (Rpl13a), ribosome biogenesis protein RLP24 (Rsl24d1), ribosome-binding protein 1 (Rbp1), release factor Gspt1, and regulatory elements, such as Igf2bp1, Gcn1l1, Larp, Fam129a and Nck1, are further indicative of the host cell translation machinery near sites of viral RNA synthesis. Notably, our results are in line with a recent genome-wide siRNA screen where translation factors were suggested to play a role in the replication of avian infectious bronchitis coronavirus (IBV) (27). The implication of this finding has, to our knowledge, not been further investigated.

Lastly, the category “catabolic processes” (Fig. 3a, S2b) includes a subset of autophagy-related factors and numerous ubiquitin-dependent ERAD components, including the E3 ubiquitin-protein ligase complex and 26S proteasome regulatory subunits (Psmc2, Psmd4). Interestingly, the importance of the ERAD pathway has also been reported in the genome-wide CRISPR screen on flaviviruses, and the ubiquitin-proteasome pathway has been noted to be important for IBV and MHV replication (27, 52).

Collectively, the catalogue of coronavirus RTC-proximal proteins greatly expands the repertoire of candidate proteins implicated in the coronavirus replication cycle, and contains several factors that have previously been reported to impact the replication of other positive-sense RNA viruses. Importantly, since our screening approach was tailored to detect host factors associated with the coronavirus RTC, it provides a spatial link of these factors to the site of viral RNA synthesis.

### Identification of proviral factors within the coronavirus RTC microenvironment

In order to assess the potential functional relevance of RTC-proximal factors identified in our MHV-BirA_R118G_-nsp2-mediated proximity-dependent screen, we designed a custom siRNA library individually targeting the expression of each of the 513 identified RTC-proximal host proteins. siRNA-treated L929 cells were infected (MOI=0.05, n=4) with a recombinant MHV expressing a *Gaussia* luciferase reporter protein (MHV-Gluc) (53) and replication was assessed by virus-mediated *Gaussia* luciferase expression (Fig. 4a). Cell viability after siRNA knockdown was also assessed and genes resulting in cytotoxicity following silencing were discarded from further analysis. Importantly, we included internal controls of known relevance for MHV entry (MHV receptor Ceacam1a) and replication (Gbf1, Arf1) on each plate and found in each case that siRNA silencing of these factors significantly reduced MHV replication, which underscores the robustness and effectiveness of our approach (Fig. S3a) (24). We found that siRNA-mediated silencing of 53 RTC-proximal host factors significantly reduced MHV replication compared to non-targeting siRNA controls. These factors can therefore be considered proviral and required for efficient replication (Fig. 3b; table S3).

**Figure 4.**
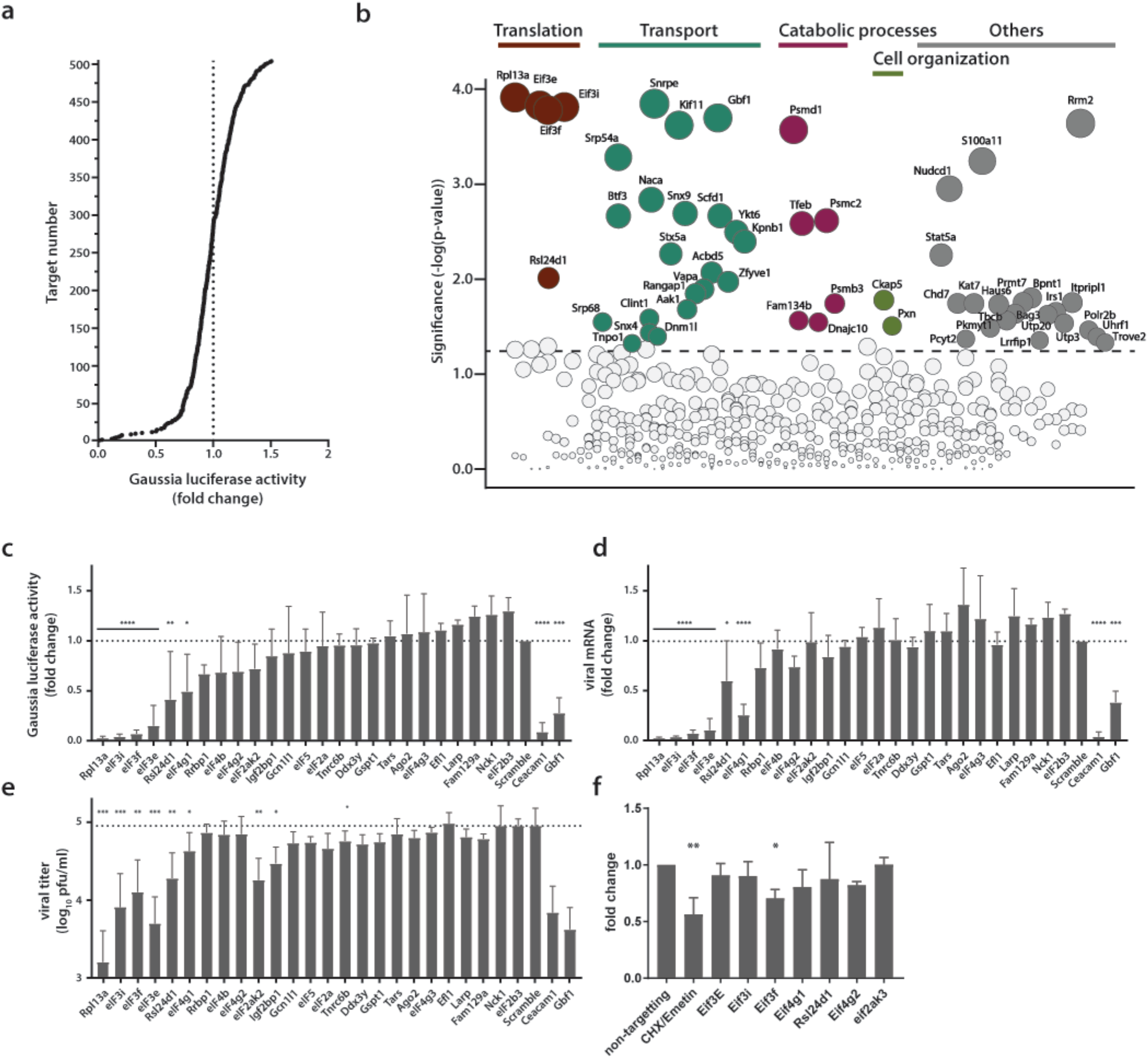
Identification of proviral factors within the coronavirus RTC microenvironment. (a) Impact of siRNA-silencing of RTC-proximal cellular proteins on viral replication. L929 fibroblasts were reverse-transfected with siRNAs (10 nM) for 48 h before being infected with MHV-Gluc (MOI=0.05, n=4). Replication was assessed by virus-mediated Gaussia luciferase expression at 15 h.p.i. and was normalized to levels of viral replication in cells targeted by scrambled siRNA controls. Target proteins to the left of the dashed line represent RTC-proximal factors whose silencing decreased viral replication. (b) Bubble plot illustrating host proteins that significantly impact MHV replication. Bubble size is proportional to the level of viral replication impairment. Colors correspond to the functional categories highlighted in Figure 3. Light grey bubbles (below the dashed line) represent host proteins that did not significantly impact MHV replication (p-value > 0.05). (c, d, e, f) Silencing of RTC-proximal components of the cellular translation machinery. Upon 48h siRNA silencing of factors assigned to the category “translation” (Figure 3), L929 fibroblasts were infected with MHV-Gluc (MOI=0.05, n=3). Luciferase activity (c), cell-associated viral RNA levels (d) and viral titers (e) were assessed at 12 h.p.i.. (f) Western blot quantification of total cellular translation following silencing of a subset of the host translation apparatus. Upon 48h siRNA-silencing, L929 fibroblasts were pulsed with 3 µM puromycin for 60 min. Control cells were treated, prior to puromycin incubation, with 355 µM cycloheximide and 208 µM Emetin for 30 min to block protein synthesis. Cell lysates were separated by SDS-PAGE and Western blots were probed using anti-puromycin antibodies to assess puromycin incorporation into polypeptides and normalized to actin levels. Error bars represent the mean ± standard deviation, where * is p ≤ 0.05, ** is p ≤ 0.005, *** is p ≤ 0.0005 and **** is p < 0.0001.

Notably, siRNA targets that had the strongest impact on MHV replication were in majority contained within the functional categories highlighted in Fig. 3a (Fig. 4b). Indeed, in line with the hypothesis that MHV subverts key components mediating both anterograde and retrograde vesicular transport between the ER, Golgi apparatus and endosomal compartments for the establishment of replication organelles, several factors contained within these pathways impaired MHV replication as exemplified by the siRNA-mediated silencing of Kif11, Snx9, Dnm11, Scfd1, Ykt6, Stx5a, Clint1, Aak11, or Vapa (Fig. 4b). Consistently, ER-associated protein sorting complexes associated with the ribosome and newly synthetized proteins (Naca, BTF3, SRP54a, SRP68) that were revealed in the GO enrichment analysis (Fig. 3a, table S2), also appear to be required for efficient MHV replication (Fig. 4b).

Furthermore, we also observed significantly reduced MHV replication upon silencing of core elements of the 26S and 20S proteasome complex (Psmd1 and Psmc2, and Psmb3, respectively), suggesting a crucial role of the ubiquitin-proteasome pathway for efficient CoV replication (27, 52). Indeed, this finding may provide a link to the described coronavirus RTC-encoded de-ubiquitination activity residing in nsp3 that has been implicated in innate immune evasion (16, 17, 54).

Most interestingly, this custom siRNA screen identified a crucial role of the host protein synthesis apparatus that was associated with the MHV RTC as indicated by the proximity-dependent proteomic screen (Fig. 3a, c). Silencing of ribosomal proteins Rpl13a and Rls24d1 and several subunits of the eIF3 complex resulted in greatly reduced MHV replication and scored with highest significance in the siRNA screen, suggesting that proximity of the host cell translation machinery to the viral RTC likely has functional importance for coronavirus replication (Fig. 4b).

### Active translation near sites of viral mRNA synthesis

Due to the striking dependence of MHV replication on a subset of RTC-proximal translation initiation factors, we extended these results in independent assays. For this, we selected all host factors assigned to the category “translation” (Fig. 3a) and assessed virus replication following siRNA-mediated silencing of each factor. Measurement of luciferase activity after MHV-Gluc infection confirmed initial findings obtained by screening the entire siRNA library of MHV RTC-proximal factors (Fig. 4c). Specifically for Rpl13a, and eIFs 3i, 3f, and 3e viral replication was reduced to levels comparable to our controls Ceacam1a (MHV receptor) and Gbf1 (24). Consistently, cell-associated viral mRNA levels (Fig. 4d) and viral titers (Fig. 4e) were reduced upon siRNA silencing of these factors. Although the silencing of a subset of host translation factors severely restricted MHV replication, effective knockdown of these factors (Fig. S3c) did not affect cell viability (Fig. S3b, d) and only moderately affected host cell translation levels (Fig. 4f, S3e). This data demonstrates that the reduced viral replication observed after siRNA knockdown is not due to a general impairment of host translation.

Subsequently, we aimed to visualize the localization of active translation during virus infection by puromycin incorporation into nascent polypeptides on immobilized ribosomes (ribopuromycylation) followed by fluorescence imaging using antibodies directed against puromycin (55). In non-infected L929 cells, ribopuromycylation resulted in an expected diffuse, mainly cytosolic, staining pattern interspersed with punctate structures indicative of translation localized to dedicated subcellular cytosolic locations (Fig. 5). In striking contrast, MHV-infected L929 cells displayed a pronounced enrichment of actively translating ribosomes near the viral RTC as indicated by the strong overlap between the viral replicase and the ribopuromycylation stain. Interestingly, active translation in vicinity of the RTC was strongest during the early phase of infection at 6 h.p.i., and was observed until 8 h.p.i., before gradually decreasing as the infection advanced along with the appearance of typical syncytia formation indicative of cytopathic effect (CPE).

**Figure 5.**
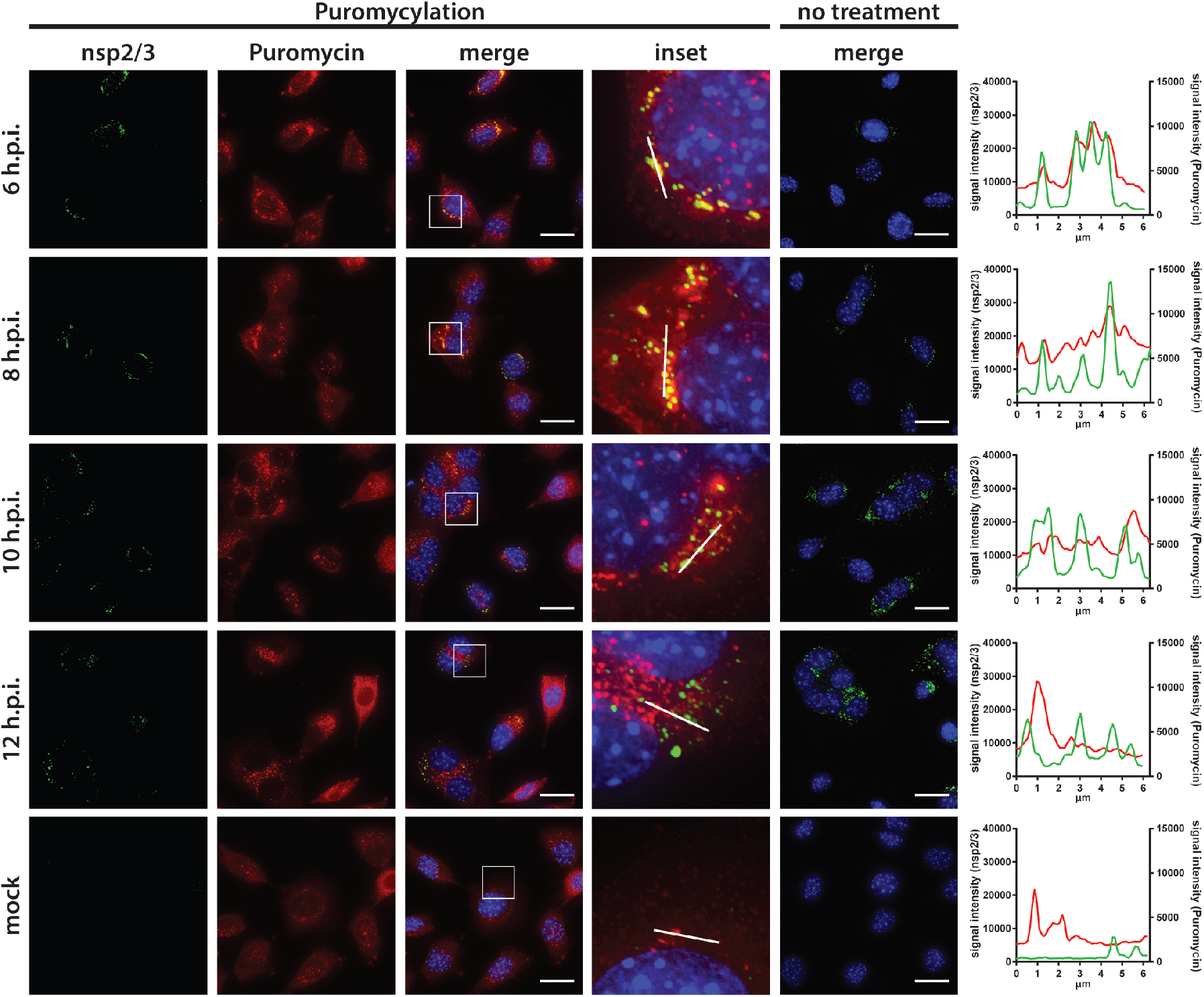
Active translation near sites of MHV mRNA synthesis. Visualization of active translation in MHV-infected L929 fibroblasts. Cells infected with MHV-A59 (MOI=1) or non-infected cells were cultured for 6, 8, 10 and 12 hours and pulsed with cycloheximide, emetine and puromycin for 5 min to label translating ribosomes. All cells, including non-treated control infections, were subjected to a coextraction/fixation procedure to remove free puromycin. Cells were labelled using anti-nsp2/3 antiserum and anti-puromycin antibodies. Nuclei are counterstained with DAPI. Z-projection of deconvolved z-stacks acquired with a DeltaVision Elite High-Resolution imaging system are shown. Note the gradual decrease of overlap between the viral replication and actively translating ribosomes highlighted in the intensity profiles. Scale bar: 20 µm.

Remarkably, we observed a similar phenotype in Huh7 cells infected with human coronaviruses, such as HCoV-229E or the highly pathogenic MERS-CoV (Fig. 6). The HCoV-229E RTC, which was detected with an antiserum directed against nsp8, appeared as small and dispersed perinuclear puncta during early infection and eventually converged into larger perinuclear structures later in infection. Consistent with findings obtained for MHV, we observed a striking co-localization of the HCoV-229E RTC with sites of active translation during the early phase of the infection (Fig. 6, S4). The co-localization gradually decreased as the infection reached the late phase with upcoming signs of CPE. Finally, we further demonstrated that active translation is localized to the site of MERS-CoV RNA synthesis as dsRNA puncta highly overlapped with the ribopuromycylation stain in MERS-CoV-infected Huh7 cells (Fig. 6). Collectively, these results not only confirm the spatial link between individual components of the host cell translation machinery and coronavirus replication compartments as identified by proximity-dependent biotinylation using MHV-BirA_R118G_-nsp2, but they also demonstrate that active translation is taking place in close proximity to the viral RTC.

**Figure 6.**
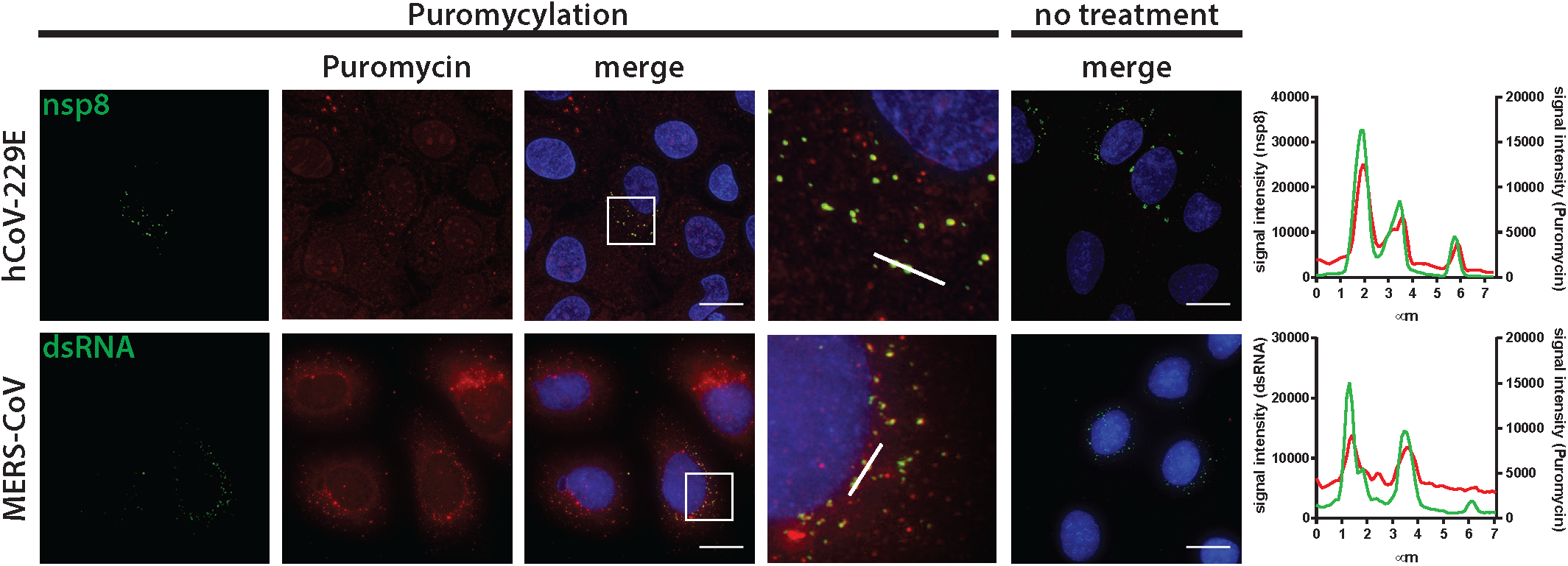
Active translation near sites of HCoV-229E and MERS-CoV mRNA synthesis. Visualization of active translation during hCoV-229E and MERS-CoV infections. Huh7 cells were infected with HCoV-229E and MERS-CoV (MOI=1) for 12 h and 6 h, respectively. Cells were pulsed with cycloheximide, emetine and puromycin for 5min to label translating ribosomes and subjected to a coextraction/fixation procedure to remove free puromycin. Non-infected and/or non-pulsed cells were used as control. Cells were labelled using anti-nsp8 (HCoV-229E) or dsRNA (MERS-CoV) and anti-puromycin antibodies. Nuclei are counterstained with DAPI. Z-projection of deconvolved z-stacks acquired with a DeltaVision Elite High-Resolution imaging system are shown. Intensity profiles of magnified regions are shown. Scale bar: 20 µm.

## Discussion

In this study, we made use of a recently developed system based on proximity-dependent biotinylation of host factors in living cells (31). By engineering a promiscuous biotin ligase (BirA_R118G_) as an integral component of the coronavirus replication complex, we provide a novel approach to define the molecular mircoenvironment of viral replication complexes that is applicable to many other RNA and DNA viruses.

We show that nsp2 fusion proteins encoded by recombinant MHV-APEX2-nsp2 and MHV-BirA_R118G_-nsp2, are indeed part of the RTC and localize to characteristic coronavirus replicative structures. On the ultrastrurctural level, APEX2-catalized DAB polymer depositions were detected at DMVs and CMs, and we observed co-localization of BirA_R118G_ with established coronavirus RTC markers, such as nsp2/3 and nsp8, by indirect immunofluorescence microscopy. Notably, in MHV-BirA_R118G_-nsp2-infected cells the detection of biotinylated coronavirus replicase gene products nsp2-10, nsp12-16, and the nucleocapsid protein by mass spectrometry demonstrates that these proteins are in close proximity during infection. This extends previous immunofluorescence and electron microscopic studies that were limited by the availability of nsp-specific antibodies and could only show localization of individual nsps to coronavirus replicative structures (4, 8, 35-37). Moreover, the close proximity of BirA_R118G_-nsp2 to MHV replicative enzymes, such as the RNA-dependent RNA polymerase (nsp12), the NTPase/helicase (nsp13), the 5’-cap methyltransferases (nsp14, nsp16), the proof-reading exonuclease (nsp14), in MHV-BirA_R118G_-nsp2-infected cells further suggest close proximity of nsp2 to the site of viral RNA synthesis. We thus propose that nsp2-16 and the nucleocapsid protein collectively constitute a functional coronavirus replication and transcription complex in infected cells.

The analysis of the host proteome enriched at MHV replication sites revealed a comprehensive list of host proteins that constitute the coronavirus RTC microenvironment. This included several individual factors and host cell pathways, especially transport mechanisms involving vesicle-mediated trafficking, which have previously been shown to assist coronavirus replication. (24, 26, 27, 49, 52). Moreover, numerous coronavirus RTC-proximal host proteins and pathways also have documented roles in the life cycle of other, more intensively studied, positive-stranded RNA viruses, suggesting considerable commonalities and conserved virus-host interactions at the replication complexes of a broad range of RNA viruses (29, 30, 56). Importantly, our list of RTC-proximal proteins by far exceeds the number of host cell proteins currently known to interact with viral replication complexes and the vast majority of MHV RTC-proximal proteins have not been described before. These likely include proteins with defined temporal roles during particular phases of the viral life cycle and proteins that did not yet attract our attention in previous screens because of functional redundancies. We therefore expect that this approach will find wide application in the field of virus-host interaction, target identification for virus inhibition, and provides a starting point to reveal similarities and differences between replication strategies of a broad range of viruses.

One novel finding that arose immediately from our RTC-proximity screen is the demonstration of a close spatial association of host cell translation with the coronavirus RTC. Indeed, the biotin ligase-based proteomic screen identified a number of translation initiation factors, most prominently several eIF3 subunits that were found to have functional importance for viral replication, and numerous ribosome-and translation-associated proteins within the coronavirus RTC microenvironment (Fig. 3, 4). In addition, the presence of subunits of the signal recognition particle in proximity to the coronavirus RTC and their functional relevance for viral replication is indicative of an importance for the translation of membrane proteins. Notably, the coronavirus RTC is translated as two polyproteins that contain nsp3, 4 and 6 with multiple trans-membrane domains that are believed to anchor the RTC at ER-derived membranes (4, 47). It is thus tempting to speculate that the coronavirus RTC is either attracting, or deliberately forming in proximity to, the ER-localized host translation machinery in order to facilitate replicase translation and insertion into ER membranes. This idea is also applicable to many other positive-stranded RNA viruses that express viral polyproteins with embedded trans-membrane domains to anchor the viral replication complex in host endomembranes. Recent experimental evidence for Dengue virus supports this hypothesis. By using cell fractionation and ribosomal profiling it has been shown that translation of the Dengue virus (family *Flaviviridae*) genome is associated with the ER-associated translation machinery accompanied by ER-compartment-specific remodeling of translation (57). Moreover, several recent genome-wide CRISPR screens demonstrated the functional importance of proteins involved in biosynthesis of membrane and secretory proteins, further supporting a pivotal role of the ER-associated translation machinery for virus replication (30).

Compartmentalization of cellular translation to sites of viral RNA synthesis has been described for dsRNA viruses of the orthoreovirus family, which replicate and assemble in distinct cytosolic inclusions known as viral factories to which the host translation machinery is recruited (58). The data presented here indicate that coronaviruses have evolved a similar strategy by compartmentalizing and directing viral RNA synthesis to sites of ER-associated translation. Likewise, this strategy has a number of advantages. Coronaviruses would not require sophisticated transport mechanisms that direct viral mRNA to distantly located ribosomes. A close spatial association of viral RNA synthesis and translation during early post-entry events would rather allow for remodeling the ER-associated translation machinery to ensure translation of viral mRNA in a protected microenvironment. Viruses have evolved diverse mechanisms to facilitate translation of their mRNAs including highly diverse internal ribosomal entry sites, recruitment of translation-associated host factors to viral RNAs, and even transcript-specific translation (59, 60). Accordingly, by remodeling defined sites for viral mRNA translation, the repertoire and concentration of translation factors can be restricted to factors needed for translation of these viral mRNAs. A microenvironment that is tailored towards the translational needs of viral mRNAs in proximity to the viral replicase complex would also make virus replication tolerant to host-or virus-induced shut down of translation at distal sites within the cytosol. Finally, proximity of viral mRNA synthesis and translation can also be considered a mechanism to evade cytosolic mRNA decay mechanisms and innate immune sensors of viral RNA.

The novel finding of a close association of the host translation machinery with sites of viral RNA synthesis during coronavirus infection exemplifies the power of the MHV-BirA_R118G_-nsp2 –mediated labelling approach to identify RTC-proximal cellular processes that significantly contribute to viral replication. Indeed, the ability of BirA_R118G_ to label viral and host factors independently of high affinity and prolonged molecular interactions enables the establishment of a comprehensive repertoire reflecting the history of protein association with the viral RTC, recorded during the entire course of infection. In future studies it will be important to provide an “RTC-association map” with temporal resolution. Like we have seen for translation initiation factors in this study, association of host cell proteins with the viral RTC might not persist throughout the entire replication cycle but might be of importance only transiently or during specific phases of the replication cycle. Given its short labelling time, APEX2 indeed offers this possibility to dissect protein recruitment to the viral RTC in a time-resolved manner, i.e. to detect RTC-associated host proteins at specific time points post infection. This will ultimately result in a dynamic, high resolution molecular landscape of virus-host interactions at the RTC and provide an additional impetus to elucidate critical virus-host interactions that take place at the site of viral RNA synthesis. These interactions should be exploited in the development of novel strategies to combat virus infection, based on conserved mechanisms of interactions at replication complexes of a broad range of positive-stranded RNA viruses.

## Acknowledgements

We thank Mark Denison, Susan Baker, and John Ziebuhr for sharing virus sequence information and antisera. We thank Sandra Huber and Kerry Woods for helpful discussions.

This work was supported by the Swiss National Science Foundation (SNF; grants # 310030_173085, and # CRSII3_160780 to V.T.). S.P. was supported by the European Commission’s Horizon 2020 research and innovation program under the Marie Sklodowska-Curie grant agreement no. 748627.

## Author Contributions

Conceptualization, P.V., V.T.; Investigation, P.V., M.G., S.P., N.E., S.B.L., J.P., H.P., V.G., R.D.; Formal Analysis, J.K., S.B.L., C.S., M.H.; Writing – Original Draft, P.V., V.T.; Supervision, V.T., R.D., M.H., R.B., M.S.; Funding Acquisition, V.T., R.D., M.H., R.B., M.S., S.P.

## Declaration of Interests

The authors declare no competing interests

## Methods

### Cells

Murine L929 fibroblasts (Sigma) and murine 17Cl1 fibroblasts (gift from S.G. Sawicki) were cultured in MEM supplemented with 10% (v/v) heat-inactivated fetal bovine serum (FBS), 100 μg/ml streptomycin and 100 IU/ml penicillin (MEM+/+). Huh-7 hepatocarcinoma cells (gift from V. Lohnmann) and Vero B4 cells (kindly provided by M. Müller) were propagated in Dulbecco’s Modified Eagle Medium-GlutaMAX supplemented with, 1 mM sodium pyruvate, 10% (v/v) heat-inactivated fetal bovine serum, 100 μg/ml streptomycin, 100 IU/ml penicillin and 1% (w/v) non-essential amino acids.

### Viruses

Recombinant MHV strain A59 (WT), MHV-Gluc (61), which expresses a *Gaussia* luciferase reporter replacing accessory gene 4 of MHV strain A59, and HCoV-229E were generated as previously described(32, 33, 62). Viruses were propagated on 17Cl1 cells (MHV) and Huh-7 cells (hCoV-229E) and their sequence was confirmed by RT-PCR sequencing. MERS-CoV (63, 64) was propagated and titrated on Vero cells.

### Generation of recombinant MHV viruses

Recombinant MHV viruses were generated using a vaccinia virus-based system as described before (33). In short, a pGPT-1 plasmid encoding an *Escherichia coli* guanine phosphoribosyltransferase (GPT) flanked by MHV-A59 nt 447-950 and 1315-1774 was used for targeted homologous recombination with a vaccinia virus (VV) containing a full-length cDNA copy of the MHV-A59 genome (32). The resulting GPT-positive VV was further used for recombination with a plasmid containing the EGFP coding sequence flanked by MHV-A59 nt 477-956 and 951-1774 for the generation of MHV-GFP-nsp2, based on the strategy employed by Freeman et al.(34). Alternatively, a plasmid containing the BirA_R118G_ coding sequence (31) or the APEX2 coding sequence (65), with a N-terminal myc-tag or V5-tag, respectively, and a C-terminal (SGG)_3_ flexible linker flanked by MHV-A59 nt 477-956 and 951-1774 was used for the generation of MHV-BirA_R118G_-nsp2 and MHV-APEX2-nsp2. The resulting VV were used to generate full-length cDNA genomic fragments by restriction digestion of the VV backbone. Rescue of MHV-GFP-nsp2, MHV-BirA_R118G_-nsp2 and MHV-APEX2-nsp2 was performed by electroporation of capped *in vitro* transcribed recombinant genomes into a BHK-21-derived cell line stably expressing the nucleocapsid (N) protein layered on permissive 17Cl1 mouse fibroblasts. Recombinant MHV viruses were plaque-purified three times and purified viruses were passaged three times for stock preparations. All plasmid sequences, VV sequences and recombinant MHV sequences were confirmed by PCR or RT-PCR sequencing. Viruses were propagated on 17Cl1 cells and virus stocks were titrated by plaque assay on L929 cells.

### Viral replication assay

L929 cells were infected with MHV-A59, MHV-GFP-nsp2, MHV-BirA_R118G_-nsp2 or MHV-APEX2-nsp2 in quadruplicate at an MOI=1. Virus inoculum was removed 2 h.p.i., cells were washed with PBS and fresh medium was added. Viral supernatants were collected at the indicated time point and titrated by plaque assay on L929 cells. Titers reported are the averages of three independent experiments ± standard error of the mean (SEM).

### Immunofluorescence imaging

Biotinylation assays were carried out as described before with minor modifications(66). 10^6^ L929 cells grown on glass coverslips were infected with MHV-A59, MHV-BirA_R118G_-nsp2 or MHV-APEX2-nsp2 at an MOI=1, or non-infected in medium supplemented with 67 µM biotin (Sigma B4501). Cells were washed three times with PBS at the indicated time points and fixed with 4% (v/v) neutral buffered formalin before being washed three additional times. Cells were permeabilized in PBS supplemented with 50 mM NH_4_Cl, 0.1% (w/v) Saponin and 2% (w/v) BSA (CB) for 60 min and incubated 60 min with the indicated primary antibodies diluted in CB (polyclonal anti-MHV-nsp2/3 or nsp8 (gift from S. Baker), 1:200 (35, 67); anti-myc, 1:8000 Cell Signalling 2276). Cells were washed three times with CB and incubated for 60 min with donkey-derived, AlexaFluor488-conjugated anti-mouse IgG (H+L) and donkey-derived, AlexaFluor647-conjugated anti-rabbit IgG (H+L) (Jackson Immunoresearch). Cells were additionally labelled with streptavidin conjugated to AlexaFluor 594 (Molecular Probes) to detect biotinylated proteins. Coverslips were mounted on slides using ProLong Diamond Antifade mountant containing 4’,6-diamidino-2-phenylindole (DAPI) (Thermo Fisher Scientific).

For indirect immunofluorescence detection of viral and host proteins, L929 cells were grown on glass coverslips in 24-well plates and infected with MHV-A59 or MHV-BirA_R118G_-nsp2 (MOI=1). At the indicated time point, cells were fixed with 4% (v/v) formalin and processed using primary monoclonal antibodies directed against dsRNA (J2 Mab, English Scientific and Consulting) or myc-tab (Cell signalling 2276) and polyclonal antibodies recognizing eIF3E (Sigma, HPA023973) or RTN4 (Nogo A+B, Abcam 47085) as well as secondary donkey-derived, AlexaFluor488-conjugated anti-mouse and AlexaFluor647-conjugated anti-rabbit IgG (H+L), as described above.

For proximity ligation assays, L929 cells were seeded in 24-well plates on glass coverslips and infection with MHV-A59 or MHV-BirA_R118G_-nsp2 (MOI=1). At the indicated time point, cells were washed with PBS, fixed with 4% (v/v) formalin and permeabilized with 0.1% (v/v) Triton X-100. Proximity ligation was performed as recommended by the manufacturer (Duolink In Situ detection reagents Red, Sigma) using monoclonal antibodies directed against dsRNA (J2, English & Scientific Consulting) or myc-tag (Cell Signaling 2276) and polyclonal antibodies recognizing eIF3E (Sigma, HPA023973) or RTN4 (Nogo A+B, Abcam 47085). Coverslips were mounted using Duolink^®^ In Situ Mounting Media with DAPI (Sigma).

All samples were imaged by acquiring 0.2 µm stacks over 10 µm using a DeltaVision Elite High-Resolution imaging system (GE Healthcare Life Sciences) equipped with a 60x or 100x oil immersion objective (1.4 NA). Images were deconvolved using the integrated softWoRx software and processed using Fiji (ImageJ).

### Biotinylation assay – western blot – mass spectrometry

L929 cells were infected with MHV-A59 or MHV-BirA_R118G_-nsp2, and for comparison MHV_H277A_ and MHV_H227A_-BirA_R118G_-nsp2, at an MOI=1 in medium supplemented with 67 µM biotin (Sigma B4501). At 15 h.p.i., cells were washed three times with PBS and lysed in ice-cold buffer containing 50 mM TRIS-Cl pH 7.4, 500 mM NaCl, 0.2% (w/v) SDS, 1 mM DTT and 1x protease inhibitor (cOmplete Mini, Roche). Cells were scraped off the flask and transferred to tubes. Cells were kept on ice until the end of the procedure. Triton X-100 was added to each sample to a final concentration of 2%. Samples were sonicated for two rounds of 20 pulses with a Branson Sonifier 250 (30% constant, 30% power). Equal volumes of 50 mM TRIS-Cl were added to each sample and samples were centrifuged at 4 °C for 10 min at 18000 x *g*. Supernatants were incubated with magnetic beads on a rotator at 4 °C overnight (800 µl Dynabeads per sample, MyOne Streptavidin C1, Life Technologies) that were previously washed with lysis buffer diluted 1:1 with 50 mM TRIS-Cl. Beads were washed twice with buffer 1 (2% (w/v) SDS), once with buffer 2 (0.1% (w/v) deoxycholic acid, 1% (v/v) Triton X-100, 1 mM EDTA, 500 mM NaCl, 50 mM HEPES pH 7.5), once with buffer 3 (0.5% w/v deoxycholic acis, 0.5% NP40, 1 mM EDTA, 250 mM LiCl, 10 mM TRIS-Cl pH 7.4) and once with 50 mM TRIS-Cl pH 7.4. Proteins were eluted from beads by the addition of 0.5 mM biotin and Laemmli SDS-sample buffer and heating at 95 °C for 10 min. For SDS-PAGE and western blot analysis, cells were cultured in 6-well plates and lysates were prepared and affinity purified as described above. Proteins were separated on 10% (w/v) SDS-polyacrylamide gels (Bio-Rad), and proteins were electroblotted on nitrocellulose membranes (Amersham Biosciences, GE Healthcare) in a Mini Trans-Blot cell (Bio-Rad). Membranes were incubated in a protein-free blocking buffer (Advansta) and biotinylated proteins were probed by incubation with horseradish peroxidase-conjugated Streptavidin (Dako). Proteins were visualized using WesternBright enhanced chemiluminescence horseradish peroxidase substrate (Advansta) according to the manufacturer’s protocol.

For mass spectrometry analysis, lysates and affinity purification were performed as described above from 4*10^7^ cells cultured in 150 cm^2^ tissue culture flasks. Proteins were separated 1 cm into a 10% (w/v) SDS-polyacrylamide gel. A Coomassie stain was performed and 4x 2 mm bands were cut with a scalpel. Proteins on gel samples were reduced, alkylated and digested with Trypsin(68). Digests were loaded onto a pre-column (C18 PepMap 100, 5 µm, 100 A, 300 µm i.d. x 5 mm length) at a flow rate of 20 µL/min with solvent C (0.05% TFA in water/acetonitrile 98:2). After loading, peptides were eluted in back flush mode onto the analytical Nano-column (C18, 3 μm, 100 Å, 75 μm x 150 mm, Nikkyo Technos C. Ltd., Japan) using an acetonitrile gradient of 5% to 40% solvent B (0.1% (v/v) formic acid in water/acetonitrile 4,9:95) in 40 min at a flow rate of 400 nL/min. The column effluent was directly coupled to a Fusion LUMOS mass spectrometer (Thermo Fischer, Bremen; Germany) via a nano-spray ESI source. Data acquisition was made in data dependent mode with precursor ion scans recorded in the orbitrap with resolution of 120’000 (at m/z=250) parallel to top speed fragment spectra of the most intense precursor ions in the Linear trap for a cycle time of 3 seconds maximum. Spectra interpretation was performed with Easyprot on a local, server run under Ubuntu against a forward + reverse *Mus musculus* (UniprotKB version 2016_04) and MHV (UniprotKB version 2016_07) database, using fixed modifications of carboamidomethylated on Cysteine, and variable modification of oxidation on Methionine, biotinylation on Lysine and on protein N-term, and deamidation of Glutamine and Asparagine. Parent and fragment mass tolerances were set to 10 ppm and 0.4 Da, respectively. Matches on the reversed sequence database were used to set a Z-score threshold, where 1% false discoveries (FDR) on the peptide spectrum match level had to be expected. Protein identifications were only accepted, when two unique peptides fulfilling the 1% FDR criterion were identified. MS identification of biotinylated proteins was performed in three independent biological replicates. For label-free protein quantification, LC-MS/MS data was interpreted with MaxQuant (version 1.5.4.1) using the same protein sequence databases and search parameters as for EasyProt. Match between runs was activated, however samples from different treatments were given non-consecutive fraction numbers in order to avoid over-interpretation of data. The summed and median normalized top3 peptide intensities extracted from the evidence table as a surrogate of protein abundance (69) and LFQ values were used for statistical testing. The protein groups were first cleared from all identifications, which did not have at least two valid LFQ values. Protein LFQ levels derived from MaxQuant were log-transformed. Missing values were imputed by assuming a normal distribution between sample replicates. A two-tailed t-test was used to determine significant differences in protein expression levels between sample groups and p-values were adjusted for multiple testing using the Benjamini-Hochberg (FDR) test.

### Computational analysis

Database for Annotation, Visualization, and Integrated Discovery (DAVID) was used to perform GO enrichment analysis on the RTC-proximal cellular factors identified via mass spectrometry(70-73). GO BP terms with a p-value <0.05 were considered to be terms that were significantly enriched in the dataset. Additional analysis of significant GO terms was conducted using AmiGO and revealed that the top 32 GO BP terms (p-value <0.005) were predominantly associated with five broad functional categories (cell-cell adhesion, transport, cell organization, translation, and catabolic processes)(74). Alternatively, enrichment analysis was performed using SetRank (data not shown), a recently described algorithm that circumvents pitfalls of commonly used approaches and thereby reduces the amount of false-positive hits (75) and the following databases were searched for significant gene sets: BIOCYC (76), GO (72), ITFP (77), KEGG (78), PhosphoSitePlus (79), REACTOME (80), and WikiPathways (81). Both independent approaches lead to highly similar results and consistently complement results obtained upon GO Cellular Components analysis.

STRING functional protein association networks were generated using RTC-proximal host proteins found within each of the five broad functional categories. Default settings were used for active interaction sources and a high confidence interaction score (0.700) was used to maximize the strength of data support. The MCL clustering algorithm was applied to each STRING network using an inflation parameter of 3 (82, 83).

### siRNA screen

A custom siRNA library targeting each individual RTC-proximal factor (On Target Plus, SMART pool, 96-well plate format, Dharmacon, GE Healthcare) was ordered. 10 nM siRNA were reverse transfected into L929 cells (8*10^3^ cells per well) using Viromer Green (Lipocalyx) according to the manufacturer’s protocol. Cells were incubated 48 hours at 37 °C 5% CO_2_ and cell viability was assessed using the CytoTox 96^®^ Non-Radioactive Cytotoxicity Assay (Promega). Cells were infected with MHV-Gluc (MOI=0.05, 1000 plaque forming units/well), washed with PBS 3 h.p.i. and incubated in MEM+/+ for additional 12 hours. Gaussia luciferase was measured from the supernatant using Pierce™ Gaussia Luciferase Glow Assay Kit (ThermoFisher Scientific). Experiments were carried out in 4 independent replicates and both cytotoxicity values and luciferase counts were normalized to the corresponding non-targeting scrambled control of each plate. A one-way ANOVA (Kruskal-Wallis test, uncorrected Dunn’s test) was used to test the statistical significance of reduced viral replication (mean < 95% as compared to scramble control, n=216). The R package ggplot2 was used to create the bubble plot (Fig 4B).

### siRNA screen validation

L929 cells were transfected with 10 nM siRNA as described above. 48 h post-transfection, cell viability was assessed using the CytoTox 96^®^ Non-Radioactive Cytotoxicity Assay (Promega) and visually inspected by automated phase-contrast microscopy using an EVOS FL Auto 2 Imaging System equipped with a 4x air objective. Cells were infected with MHV-Gluc (MOI=0.05), washed with PBS 3 h.p.i. and incubated for 9 additional hours. *Gaussia* luciferase activity, viral titers and cell viability were measured from the supernatant as described above. One-way ANOVAs (ordinary one-way ANOVA, uncorrected Fisher’s LSD test) were used to test the statistical significance.

Total cellular RNA was isolated from cells using the NucleoMag^®^ RNA Kit (Machery Nagel, Switzerland) on a KingFisher™ Flex Purification System (Thermo Fisher Scientific, Switzerland) according to the manufacture’s instructions. The QuantiTect Probe RT-PCR Kit (Qiagen, Switzerland) was used according to the manufactures instructions for measuring the cell associated viral RNA levels with primers and probe specific to the MHV genome fragment coding the nucleocapsid gene (Table S4). Primers and Probe for mouse Glyceraldehyde 3-phosphate dehydrogenase (GAPDH) where obtained from ThermoFisher Scientific (Mm03302249_g1, Catalog Number: 4331182). The MHV levels were normalized to GAPDH and shown as ΔΔCt over mock (ΔCt values calculated as Ct reference - Ct target). The QuantiTect SYBR^®^ Green RT-PCR Kit (Qiagen, Switzerland) was used according to the manufactures instructions for measuring the expression levels of Rpl13a, eIF3E, eIF3I, eIF3F, eIF4G1, eIF4G2, eIF2ak3, Rsl24d1 and Tbp. All primer pairs where placed over an exon intron junction (Table S4). All expression levels are displayed as ΔΔCt over non-targeting siRNA (ΔCt values calculated as Ct target - Ct Tbp) (84). One-way ANOVA (ordinary one-way ANOVA, uncorrected Fisher’s LSD test) was used to test the statistical significance.

### Total cellular translation

siRNA-based silencing was performed as described above. 48 h post-transfection, control cells were incubated with 355 μM cycloheximide (Sigma) and 208 μM Emetin (Sigma) for 30 min to block protein synthesis. Cells were treated with 3 μM puromycin for 60 min followed by three PBS washes(85). Total cell lysates were prepared using M-PER mammalian protein extraction reagent (Thermo Scientific) supplemented with protease inhibitors (cOmplete Mini, Roche). Lysates were separated on a 10% (w/v) SDS-PAGE and electroblotted as described above. Western blots were probed using a monclonal AlexaFluor647-conjugated anti-puromycin antibody (clone 12D10, Merk Millipore) and a donkey-derived HRP-conjugated anti-mouse (Jackson immunoresearch 715-035-151). Actin was detected using a monoclonal HRP-conjugated anti-actin antibody (Sigma A3854) and used to normalize input.

### Ribopuromycylation assay

Ribopuromycylation of actively translating ribosomes was performed as described before (55). L929, Huh-7 cells were seeded on glass coverslips and infected with MHV-A59 (L929), HCoV-229E (Huh-7), MERS-CoV (Huh-7) and at MOI=1. One hour after inoculation, cells were washed with PBS and incubated further for the indicated time. Cells were treated with 355 μM cycloheximide and 208 μM Emetin (Sigma) for 15 min at 37°C. Cells were further incubated in medium containing 355 μM cycloheximide, 208 μM Emetin and 182 μM puromycin (Sigma) for additional 5 min. Cells were washed twice in ice-cold PBS and fix on ice for 20 min in buffer containing 50 mM TRIS HCl, 5 mM MgCl_2_, 25 mM KCl, 355 μM cycloheximide, 200 mM NaCl, 0.1% (v/v) TritonX-100, 3% formalin and protease inhibitors (cOmplete Mini, Roche). Cells were blocked for 30 min in CB, and immunostained as described above using polyclonal anti-MHV-nsp2/3 (gift from S. Baker), polyclonal anti-HCoV-229E-nsp8 (gift from J. Ziebuhr), or monoclonal anti-dsRNA (J2 MAB, English and Scientific Consulting) as primary antibodies to detect MHV, HCoV-229E and ZIKV replication complexes, respectively. Donkey-derived, AlexaFluor488-conjugated anti-mouse or anti-rabbit IgG (H+L) were used as secondary antibodies. Additionally, ribosome-bound puromycin was detected using a monoclonal AlexaFluor647-conjugated anti-puromycin antibody (clone 12D10, Merk Millipore). Slides were mounted, imaged and processed as described above.

### DAB staining and transmission electron microscopy

L929 fibroblasts were seeded in 24-well plates and infected with MHV-APEX2-nsp2, MHV-A59, or non-infected for 10 h. 3,3-diaminobenzidine (DAB) stains were performed as described previously (38). Briefly, cells were fixed at 10 h.p.i. using warm 2% (v/v) glutaraldehyde in 100 mM sodium cacodylate, pH 7.4, supplemented with 2 mM calcium chloride (cacodylate buffer) and placed on ice for 60 min. The following incubations were performed on ice in ice-cold buffers unless stated otherwise. Cells were washed 3x with sodium cacodylate buffer, quenched with 20 mM glycine in cacodylate buffer for 5 min. before 3 additional washes with cacodylate buffer. Cells were stained in cacodylate buffer containing 0.5 mg/ml DAB and 10 mM H2O2 for 20 min until DAB precipitates were visible by light microscopy. Cells were washed 3x with cacodylate buffer to stop the staining reaction. Processing of samples for transmission electron microscopy (TEM) was performed as described previously (86). Briefly, cells were washed once with PBS prewarmed to 37 °C and subsequently fixed with 2.5% (v/v) glutaraldehyde (Merck, Darmstadt, Germany) in 0.1 M cacodylate buffer (Merck, Hohenbrunn, Germany) pH 7.4 for 30 min at room temperature or overnight at 4 °C. After three washes in cacodylate buffer for 10 min each, cells were post-fixed with 1% OsO4 (Chemie Brunschwig, Basel, Switzerland) in 0.1 M cacodylate buffer for 1 h at 4 °C and again washed three times with cacodylate buffer. Thereafter, cells were dehydrated in an ascending ethanol series (70%, 80%, 90%, 94%, 100% (v/v) for 20 min each) and embedded in Epon resin, a mixture of Epoxy embedding medium, dodecenyl succinic anhydride (DDSA) and methyl nadic anhydride (MNA) (Sigma Aldrich, Buchs, Switzerland). Ultrathin sections of 90 nm were then obtained with diamond knives (Diatome, Biel, Switzerland) on a Reichert-Jung Ultracut E (Leica, Heerbrugg, Switzerland) and collected on collodion-coated 200-mesh copper grids (Electron Microscopy Sciences, Hatfield, PA, USA). Sections were double-stained with 0.5% (w/v) uranyl acetate for 30 min at 40 °C (Sigma Aldrich, Steinheim, Germany) and 3% (w/v) lead citrate for 10 min at 20 °C (Laurylab, Saint Fons, France) in an Ultrastain^®^ (Leica, Vienna, Austria) and examined with a Philips CM12 transmission electron microscope (FEI, Eindhoven, The Netherlands) at an acceleration voltage of 80 kV. Micrographs were captured with a Mega View III camera using the iTEM software (version 5.2; Olympus Soft Imaging Solutions GmbH, Münster, Germany).

## Figure legends

**Supplemental Figure 1.**
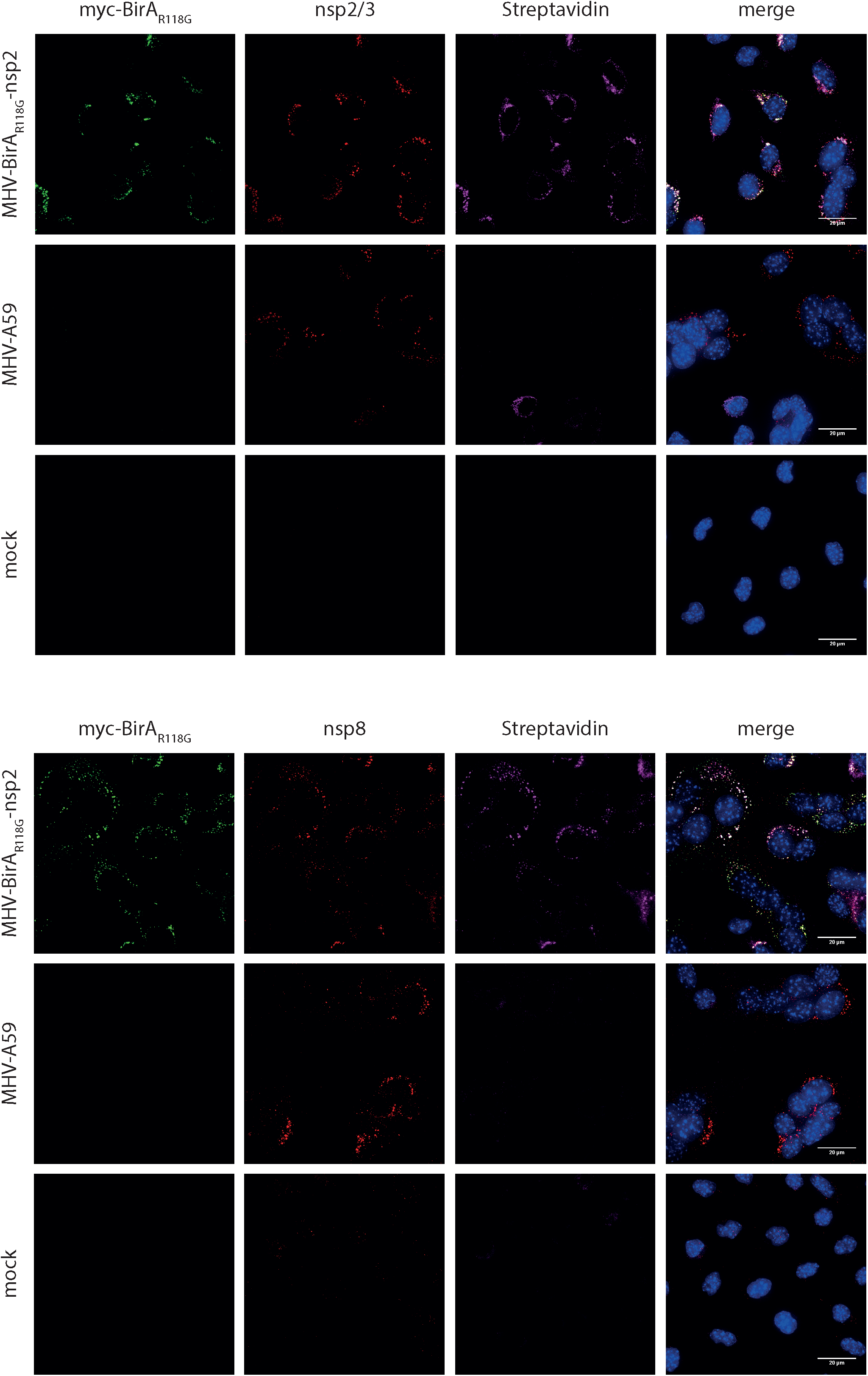
Immunofluorescence analysis of MHV-BirA_R118G_-nsp2-mediated biotinylation. MHV-BirA_R118G_-nsp2, MHV-A59-or non-infected L929 fibroblasts were cultured in medium supplemented with 67µM biotin. Cells were fixed 12 hours post infection (h.p.i.) and processed for immunofluorescence analysis with antibodies directed against the BirA_R118G_ (anti-myc), the viral replicase (anti-nsp2/3 or nsp8) and biotinylated factors (streptavidin). Nuclei are counterstained with DAPI. Z-projection of deconvolved z-stacks acquired with a DeltaVision Elite High-Resolution imaging system are shown. Scale bars: 20 µm.

**Supplemental Figure 2.**
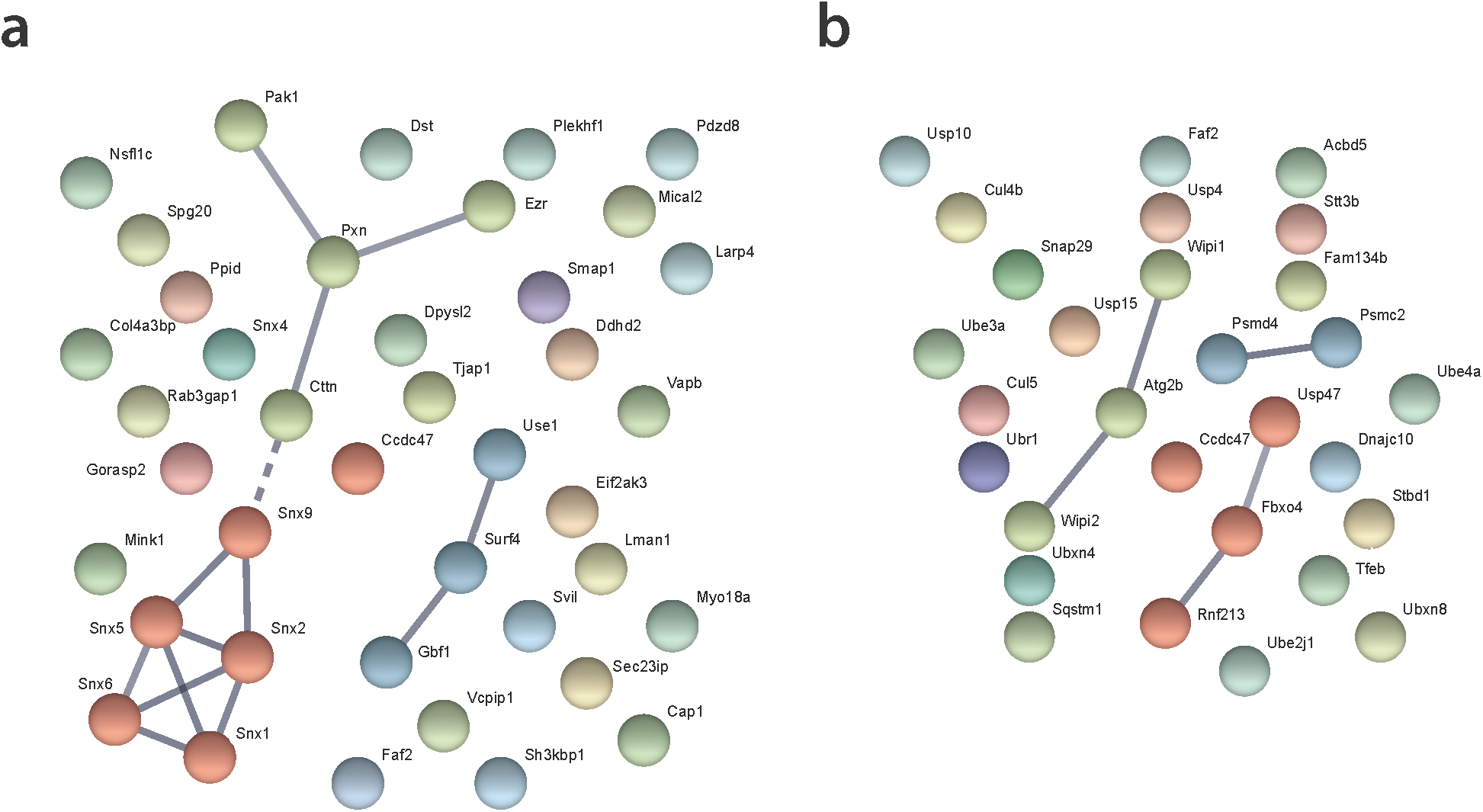
STRING protein interaction network analysis of the categories “cell organization” (a) and “catabolic processes” (b). The nodes represent RTC-proximal host proteins and the edges represent the interactions, either direct (physical) or indirect (functional), between two proteins in the network.

**Supplemental Figure 3.**
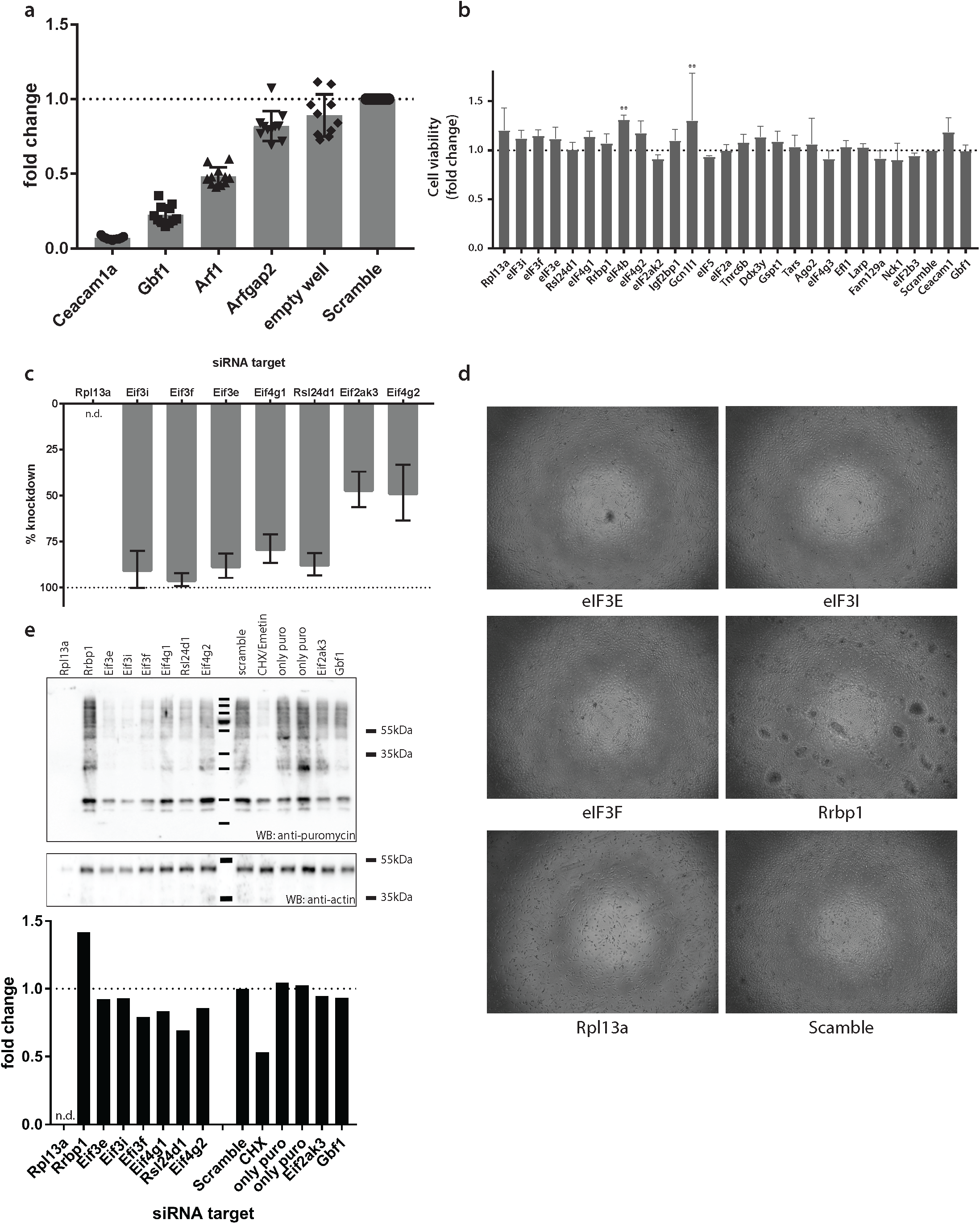
(a) siRNA controls contained in each 96-well plate during siRNA-silencing of the RTC-proximal library. Controls included the established factors such as MHV entry receptor (Ceacam1a), Gbf1, Arf1. Arfgap2 was found to moderately affect MHV replication during pilot experiments and was included to cover the entire inhibitory range. (b) Cell viability following 48h siRNA-silencing of components of the cellular translation machinery. (c) Expression levels of Rpl13a, eIF3E, eIF3I, eIF3F, eIF4G1, eIF4G2, eIF2ak3, Rsl24d1 following siRNA knockdown compared to expression levels in cells treated with non-targetting siRNA. (d) Visual inspection of L929 treated with siRNA targetting eIF3E, eIF3I, eIF3F, Rrbp1, Rpl13a, non-targetting siRNA (scramble). Note that RNA silencing (b) and translation activity (c) in Rpl13a-silenced cells could not be assessed, likely due to cytotoxicity observed by visual inspection of cells. (e) Western blot and western blot analysis of total cellular translation. Upon 48h siRNA-silencing, L929 fibroblasts were pulsed with 3 µM puromycin for 60 min. Control cells were treated, prior to puromycin incubation, with 355 µM cycloheximide and 208 µM Emetin for 30 min to block protein synthesis. Western blots were probed using anti-puromycin antibodies to assess puromycin incorporation into polypeptides and normalized to actin levels. Error bars represent the mean ± standard deviation of three independent experiments, where * is ** is p ≤ 0.005.

**Supplemental Figure 4.**
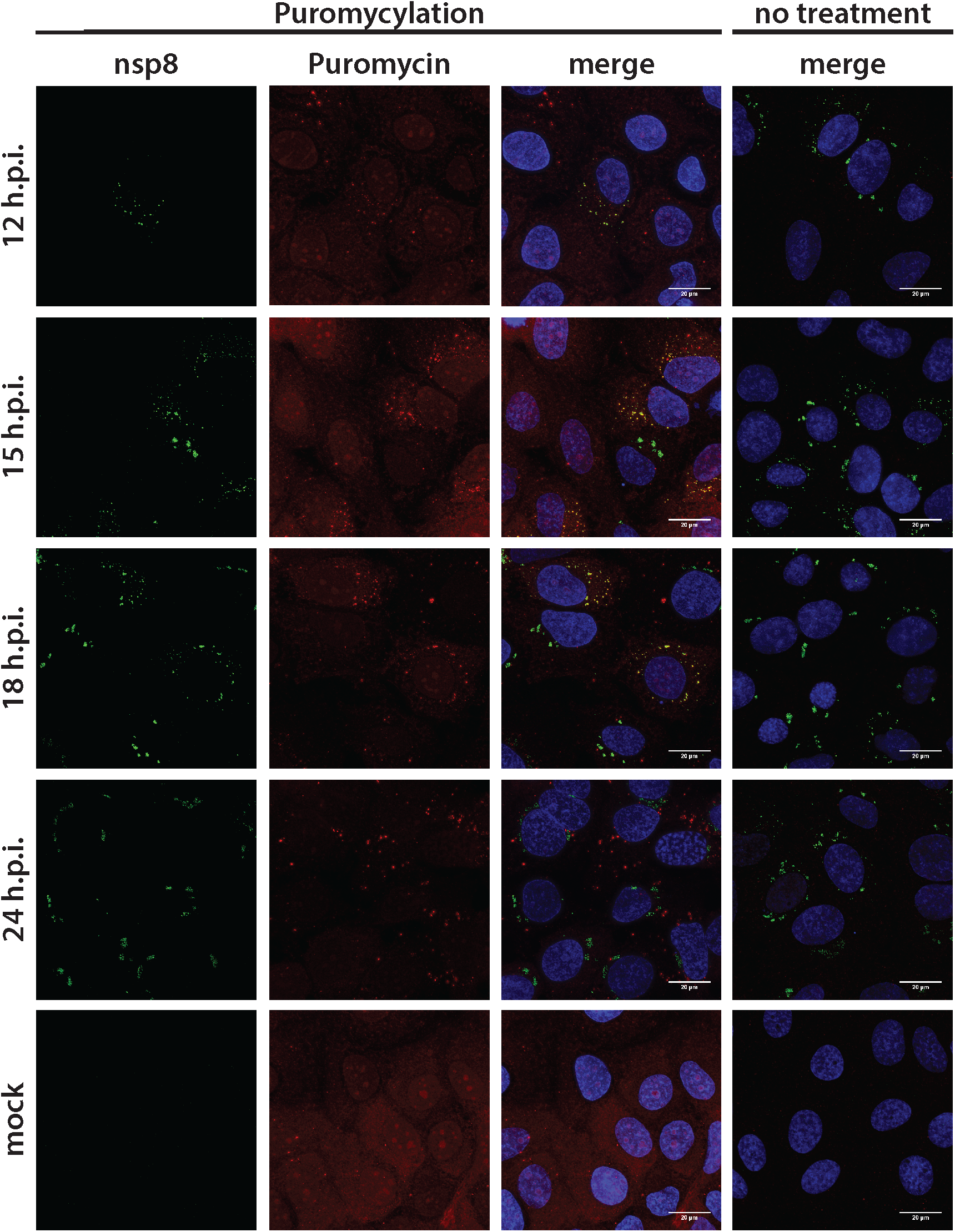
Visualization of active translation during HCoV-229E infections. Huh7 cells were infected with HCoV-229E (MOI=1) for 12, 15, 18, 24 h. Cells were pulsed with cycloheximide, emetine and puromycin for 5min to label translating ribosomes and subjected to a coextraction/fixation procedure to remove free puromycin. Non-infected and/or non-pulsed cells were used as control. Cells were labelled using anti-nsp8 (HCoV-229E) and anti-puromycin antibodies. Nuclei are counterstained with DAPI. Z-projection of deconvolved z-stacks acquired with a DeltaVision Elite High-Resolution imaging system are shown. Intensity profiles of magnified regions are shown. Scale bar: 20 µm.

